# Repair of critical-size porcine craniofacial bone defects using a collagen-polycaprolactone composite biomaterial

**DOI:** 10.1101/2021.04.19.440506

**Authors:** Marley J. Dewey, Derek J. Milner, Daniel Weisgerber, Colleen L. Flanagan, Marcello Rubessa, Sammi Lotti, Kathryn M. Polkoff, Sarah Crotts, Scott J. Hollister, Matthew B. Wheeler, Brendan A.C. Harley

## Abstract

Regenerative medicine approaches for massive craniomaxillofacial bone defects face challenges associated with the scale of missing bone, the need for rapid graft-defect integration, and challenges related to inflammation and infection. Mineralized collagen scaffolds have been shown to promote mesenchymal stem cell osteogenesis due to their porous nature and material properties, but are mechanically weak, limiting surgical practicality. Previously, these scaffolds were combined with 3D-printed polycaprolactone mesh to form a scaffold-mesh composite to increase strength and promote bone formation in sub-critical sized porcine ramus defects. Here, we compare the performance of mineralized collagen-polycaprolactone composites to the polycaprolactone mesh in a critical-sized porcine ramus defect model. While there were no differences in overall healing response between groups, our data demonstrated broadly variable metrics of healing regarding new bone infiltration and fibrous tissue formation. Abscesses were present surrounding some implants and polycaprolactone polymer was still present after 9-10 months of implantation. Overall, while there was limited successful healing, with 2 of 22 implants showed substantial levels of bone regeneration, and others demonstrating some form of new bone formation, the results suggest targeted improvements to improve repair of large animal models to more accurately represent craniomaxillofacial bone healing. Notably, strategies to increase osteogenesis throughout the implant, modulate the immune system to support repair, and employ shape-fitting tactics to avoid implant micromotion and resultant fibrosis. Improvements to the mineralized collagen scaffolds involve changes in pore size and shape to increase cell migration and osteogenesis and inclusion or delivery of factors to aid vascular ingrowth and bone regeneration.

## 1. Introduction

Craniomaxillofacial (CMF) defects represent portions of bone missing from the skull or jaw that cannot be repaired without surgical intervention. CMF injuries and defects can occur throughout the lifespan of an individual, such as during birth (cleft palate defects), during wartime (high-energy impact trauma), and from cancer and dentures (loss of bone from surgery or atrophy) [1, 2]. Current methods of repair involve the use of autografts or allografts, bone from the patient or a donor, respectively. Autografts are limited by the amount of bone able to be removed and possible morbidity at the site of removal [3, 4]. Allografts are limited by the processing method, with cellular materials removed before implantation to avoid disease transmission, which can also remove beneficial components that aid in repair [5]. Biomaterial strategies have been extensively investigated to avoid the drawbacks of allografts and autografts by providing reproducible materials with modifiable factors such as pore size, growth factors, and material types to enhance bone regeneration [6]. Biomaterials for CMF bone repair need to be carefully engineered, as effectively healing these defects is challenging due to multiple factors: size and mechanics of bone defects, multiple cell types involved, and a higher change of chronic infection and inflammation [5, 7–9].

CMF defects are often irregular in size and shape and implants must be designed to cover this unique space while also ensuring a proper fit. To ease biomaterial implantation, an implant should be patient-specific or easy to handle by surgeons to cover this defect space appropriately. The implant must also be mechanically stable as to limit motion within the defect, also known as micromotion, which can directly inhibit osseointegration and result in fibrous tissue ingrowth [10, 11]. Due to the large size of the defect, the biomaterial must promote bone formation throughout the defect area in order to completely repair the bone, being aware of the multiple cell types involved, such as osteoblasts, endothelial cells, and immune cells [12]. Finally, the immune system response can halt the formation of new bone early-on if it persists [13]. Degradation byproducts of the implant can potentially elicit a persistent inflammatory response isolated to the implant site, which may delay or inhibit wound healing and ultimately lead to implantation failure [14]. Additionally, bone defects are likely to become infected with bacteria during surgery, and common infectious bacteria such as *Staphylococcus aureus* can prevent bone formation and can resist antibiotic treatments [15–17]. These challenges all lead to a complicated environment for bone regeneration and the need for precise design of biomaterial implants to address these.

Here, we investigate a mineralized collagen composite biomaterial in a critical-sized porcine ramus defect model to evaluate its potential to repair CMF defects in humans. Mineralized collagen scaffolds are porous materials composed of type I collagen, calcium phosphate mineral, and glycosaminoglycans [18]. These scaffolds have shown promise *in vitro,* demonstrating mesenchymal and adipose stem cell osteogenic differentiation and mineral formation [18–28]. Additionally, these scaffolds seeded with mesenchymal stem cells have been shown to produce osteoprotegerin (OPG), which can inhibit osteoclasts and bone resorption [29, 30]. *In vivo* studies in rabbit models with mineralized collagen scaffolds have demonstrated bone repair without stem cell or bone morphogenic protein addition [31]. A concern of use of these scaffolds in potential clinical trials is the weak mechanical properties. To address this, the mechanical properties of these scaffolds have been improved by the addition of polycaprolactone (PCL) 3D-printed support structures, which increased the stiffness 6000-fold [32]. These mineralized collagen-PCL composites (CGCaP-PCL) also demonstrated improved bone infill in sub-critical sized (10mm dia.; 10 mm thick) porcine ramus defect models compared to the mineralized collagen and PCL support alone [33].

The goal of this study was to evaluate the healing in a critical-sized (25 mm diameter; 10 mm thick) porcine mandible defect using CGCaP-PCL composites and PCL support structures. We investigated healing by micro-CT analysis of bone infill and histological evaluation of the implants after 9-10 months of implantation. We hypothesized that the CGCaP-PCL composite would repair bone within these defects and would create more new bone than PCL structures alone, similar to observations for sub-critical sized porcine defects. We report new bone infill, the immune response, and how our results motivate future biomaterial design to improve healing. We also relate our results back to the challenges to repair, notably promoting osteogenesis within the implant, limiting micromotion, and preventing a persistent immune response or infection.

## 2. Materials and Methods

### 2.1 Fabrication of 3D-printed PCL supports

PCL supports were fabricated as previously described [32, 33]. Briefly, PCL 3D prints were fabricated via laser sintering of a PCL powder with 4 wt% hydroxyapatite at Georgia Tech. The final print shape was determined by a custom MATLAB program (MathWorks^TM^), which was a cylinder measuring 25 mm diameter and 10 mm high. This cylinder was composed of repeating unit cells with dimensions of 5 mm cubes containing 3.5 mm cylindrical pores [34, 35].

### 2.2 Fabrication of CGCaP-PCL composites

CGCaP-PCL composites were fabricated as previously described [32, 36]. Briefly, this involved lyophilizing a mineralized collagen precursor suspension with a PCL support. The mineralized collagen suspension was generated via homogenizing 1.9% w/v type I collagen (Collagen Matrix, Oakland, NJ), 0.84% w/v chondroitin-6-sulfate (Sigma-Aldrich, St. Louis, MO), and calcium hydroxide (Sigma-Aldrich) and calcium nitrate tetra-hydrate (Sigma-Aldrich) and phosphoric acid. PCL 3D-prints were placed into polysulfone molds of 25 mm diameter and 10 mm height before the homogenized mineralized collagen suspension was pipetted into the mold and filled around and inside the PCL support. CGCaP-PCL composites were created by lyophilizing this suspension and PCL support using a Genesis freeze-dryer (VisTis, Gardener, NY). The suspension was frozen at a constant cooling rate of 1°C/min until reaching −10°C for 175 min, then it was sublimated at 0°C and 200 mTorr to evaporate ice crystals and create porous mineralized collagen surrounding the PCL 3D-print [36].

### 2.3 Sterilization of composites and PCL supports

PCL supports and CGCaP-PCL composites were sterilized via a 12 hour ethylene oxide treatment with an AN74i Anprolene gas sterilizer (Andersen Sterilizers Inc., Haw River, NC). Supports and composites were placed in sterilization pouches and all following handling steps prior to the *in vivo* study were done utilizing sterile techniques.

### 2.4 Hydration and crosslinking of composites

CGCaP-PCL composites and PCL supports were hydrated prior to use *in vivo*. Briefly, samples were soaked in 100% ethanol, then soaked in multiple Phosphate Buffered Saline (PBS) washes to fully hydrate the collagen [37]. The CGCaP composites were then crosslinked via carbodiimide chemistry in a PBS solution of 1-ethyl-3-(3-dimethylaminopropyl) carbodiimide hydrochloride (EDC, Sigma-Aldrich) and N-hydroxysulfosuccinimide (NHS, Sigma-Aldrich) at a molar ration of 5:2:1 EDAC:NHS:COOH [38–41]. PCL supports were not crosslinked due to lacking functional groups present on collagen. PCL supports and CGCaP-PCL composites were then washed multiple times in sterile PBS, and finally stored in fresh PBS at 37°C and 5% CO2 before use.

### 2.5 Porcine critical-sized ramus defect model

#### Surgical implantation of constructs in the porcine jaw

All described surgical procedures, associated protocols, and facilities were approved by the University of Illinois at Urbana-Champaign Institutional Animal Care and Use Committee under protocol #15065. Before surgery, all pigs received a sedative cocktail (TARK) consisting of Telazol (tiletamine and zolazepam; Pfizer, New York, NY), Atropine (Neogen Corporation, Lexington, KY), Rompun (xylaazine; Lloyd Laboratories, Shenandoah, IA), and Ketamine (Ketaset^®^; Fort Dodge Animal Health, Fort Dodge, IA) intramuscularly, then again intravenously via the ear vein canula, as necessary. Additionally, endotracheal administration of 3–5% isoflurane was administered in oxygen as anesthesia during surgery.

The surgical approach first consisted of a single submandibular and retromandibular incision tracing the exterior contour of the mandible. The underlying superficial fat and masseter muscle were then incised and elevated. Then the periosteum was incised, exposing the posterior region of the underlying hemi-mandible. Utilizing a right-angle surgical drill with a 25 mm (outer diameter) trephine (Stryker, Portage, MI), full thickness 25 mm diameter cylindrical defects were made to the exposed region of the hemi-mandible. The wound site was irrigated with 0.9% physiological saline during drilling to prevent damage to the surrounding tissue. Self-tapping bone screws (Stryker Osteosynthesis Freiburg, Freiburg, Germany) were placed adjacent to each defect as a means of localizing the defect after harvest; the distance between the wound edge and the screw was measured and recorded to aid recovery. PCL supports and CGCaP-PCL composites were then press fit into the defects and stabilized by application of Vetbond (3-M, Minneapolis, MN).

The wound site was closed by suturing the previously described incisions in the periosteum, masseter muscle, superficial fat, and finally the skin using 3-0 Vicryl (Ethicon, Inc., San Angelo, TX) in a three-layer continuous suture. Vetbond was additionally applied to the skin layer incision as an auxiliary means of closure. Following surgery, all pigs received a treatment of an antibiotic Excede (1 mL/20 kg; Pfizer) and an analgesia Banamine S (2.2 mg/kg; Schering Plough Animal Health, Summit, NJ). Both during and after surgery each pig was monitored and temperature taken every 15 min until the pig became sternally recumbent. Physical and social behavior was evaluated twice daily for the first 2 weeks post-surgery.

#### Harvesting of bone-implant samples

Implants remained in the pigs for 9-10 months before euthanasia and harvesting of the implants and surrounding bone. After euthanasia, soft tissue was dissected away from the mandible segment, and a reciprocating saw was used to separate the mandible segment from the animal. Subsequently, using a scalpel and razor blade, additional soft tissue was cleaned away from the jaw segment to aid in identifying the area housing the implant. There were no indications of implants other than callous formation and estimates of implant location were made before trimming. Once tissue was removed to expose the relevant area, a band saw was used to isolate the implant and surrounding bony tissue for further analysis. The final trimmed bone-implant samples were approximately square in shape and were roughly 3.5 cm by 3.5 cm in dimension and approximately 5 cm in thickness. After isolation and trimming, implants were fixed in 10% buffered formalin solution (Leica diagnostics) and kept at 4°C for 48 hours, then rinsed and stored in 1X PBS prior to micro-CT analysis.

### 2.6 Evaluation of bone formation via CT and micro-CT

Computed tomography (CT) scans were taken of two live pigs at 8 and 16 weeks *in vivo*. Animals were sedated with a sedative cocktail (TARK) as described previously before live imaging. Scans were taken of the entire skull with a Lightspeed 16 slice Helical CT scanner (General Electric, Fairfield, CT) at 140 kEV and 240mA. Post sacrifice, new bone formation within scaffolds and composites of each mandible was measured using microcomputed tomography (micro-CT) via a MicroXCT-400 (Zeiss, Oberkochen, Germany). Scans utilized a 0.5x camera with variable source and detector distance, power and contrast and brightness, due to variable sample size. To quantify fill of bone and mineral intensity, a custom Matlab program was used [36], which took z-stacks of 2D micro-CT images and analyzed pixels identified as bone as a function of depth, angle, and radius (**Fig. 1**). Intensity thresholds were selected by the user. Due to variable image capture conditions and intensity threshold, results of each sample were normalized to the intensity of the sample’s surrounding bone.

**Fig. 1.**
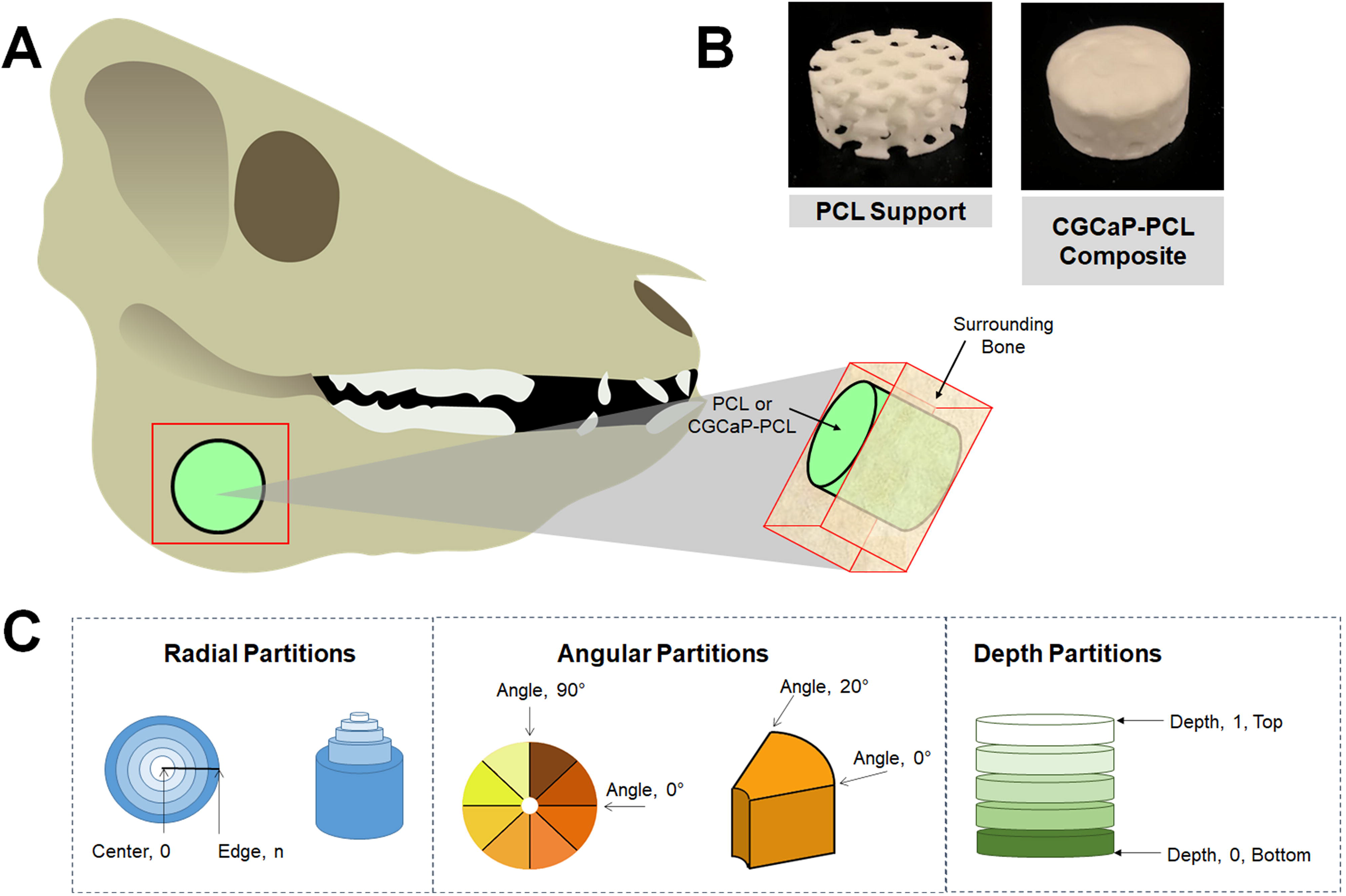
Diagram demonstrating the isolation of PCL or CGCaP-PCL composites from porcine mandible and the following Micro-CT analysis. (A) Diagram of the region of outside bone and sample of interest. A bone saw was used to cut along red region to isolate both the sample (PCL or CGCaP-PCL composite) as well as surrounding bone. (B) Images of 25 mm dia. PCL support and CGCaP-PCL composite. CGCaP completely covers in and around the PCL support. (C) Methods of analyzing sample in surrounding bone by Micro-CT, segmented by radial partitions, angular partitions, and depth partitions.

### 2.7 Histological analysis

Post micro-CT analysis, samples were rinsed again in 1X PBS, then decalcified for 4-8 days in CalciClear decalcifying solution (National Diagnostics). Samples were rinsed in several changes of water after decalcification, then overnight in 1X PBS. In order to facilitate embedding and mounting of samples for sectioning, the samples were first pre-sectioned into thick slices (~5 −10 mm) using a scroll saw, as though the decalcified bone tissue was soft enough to cut with a razor blade, the PCL support required significantly more force to cut. Thick slices were then embedded in Neg 50 cryosectioning medium (Richard Allan Scientific), and 20-30 μm sections were cut using a Leica CM1900 cryostat. Sections were then stained using hematoxylin and eosin, and images were captured using a Nanozoomer histological slide scanner (Hamamatsu) or Zeiss inverted microscope with an Axiocam color camera (Carl Zeiss).

### 2.8 Statistics

Statistical analysis used an OriginPro software (Northampton, Massachusetts) with a 95% confidence interval and testing in accordance with literature [42]. Firstly, the assumption of normality was checked using the Shapiro-Wilk test and non-normal data underwent a Grubbs outlier test to remove any outliers and normality was re-assessed. Non-normal data underwent a Kruskal-Wallis test to evaluate significant differences. If data was normal, the assumption of equal variance was then tested with a Browne-Forsythe test. If assumption was not met, a t-test with a Welch correction was used to compare two samples. A t-test was used for all analysis between only two samples. If all assumptions were met and the power was above 0.8, then an ANOVA with a tukey post-hoc test was used to determine significance. If the power was below 0.8 for any sample set, then the data was deemed inconclusive.

## 3. Results

### 3.1 Surgical results

A variety of healing outcomes were observed in the porcine models, highlighting challenges of large-scale defect models (**Table 1)**. Most notably, multiple implants were unable to be located at time of extraction, either due loss of the implant out of the wound site during healing or improper alignment during extraction. Out of 22 implants, 5 had either visibly migrated out of the jaw or were unable to be found after sacrifice. Further, PCL supports and CGCaP-PCL composites were not implanted with any location markers so extraction of the implant and surrounding bone was based on inspection of the mandible after harvest; two out of 22 implants were damaged during sample trimming due to an improper location estimate. Lastly, two out of 11 pigs were sacrificed early due to IACUC guidelines and three implants were not analyzed due to abscess formation over the implant, obscuring bone regeneration.

**Table 1.**
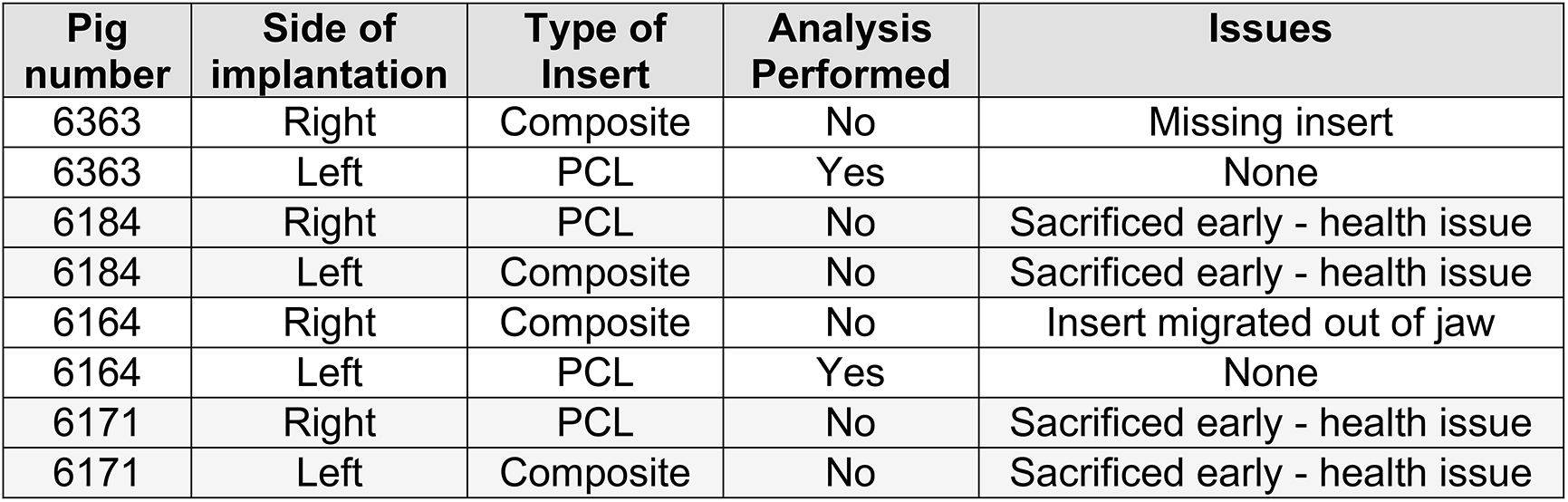
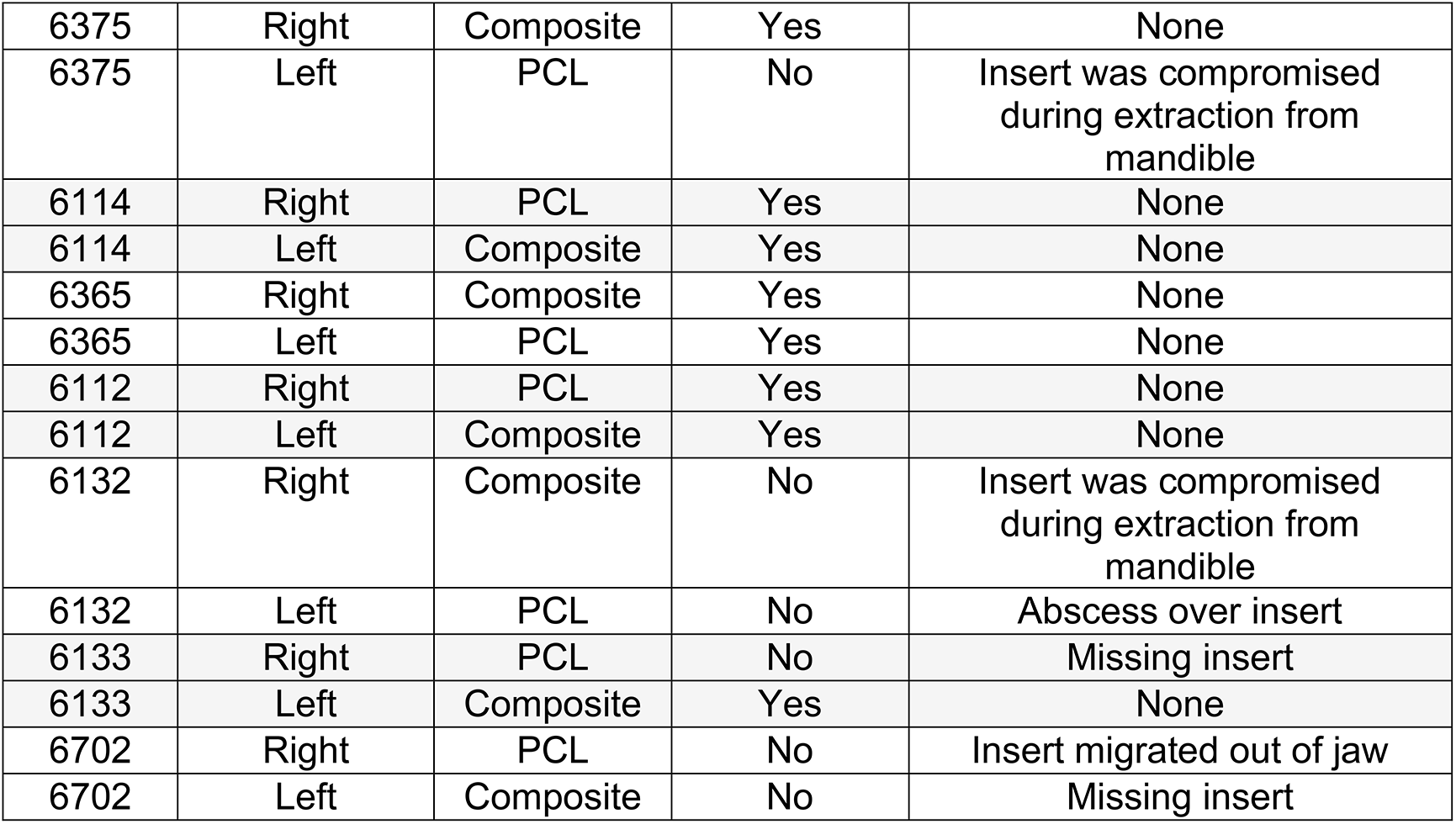
Summary of samples post-sacrifice. The following table outlines the location of each sample, PCL support or CGCaP-PCL composite (denoted as Composite) in the corresponding side of the animal jaw (right or left). Whether analysis was performed on the sample is also listed, analysis including Micro-CT and histology. Complicating issues with retrieval of the implants or with the specimen are also included.

### 3.2 No differences in new bone formation between CGCaP-PCL composites and PCL supports

Micro-CT analysis of live pigs with PCL supports and CGCaP-PCL composites was performed at 8 weeks and 16 weeks post-operation and pre-sacrifice. New bone formation is visible in both constructs, but with little to no change from 8 weeks to 16 weeks (**Supp. Fig. 1**). Micro-CT analysis of CGCaP-PCL composites and PCL supports post-sacrifice demonstrated no significant (p < 0.05) difference in fill fraction or intensity of the two implants as a function of radius, depth, or angle (**Fig. 2**). A value of 1 for the fill fraction represents the same amount of bone present as the normal surrounding bone, while a value of 1 for the intensity represents the mineral density of bone in the normal surrounding bone. Fill fraction values greater than 1 represent more bone within the implant space and intensity values greater than 1 represent a more dense and mineralized bone present. The radial fill fraction demonstrates that there was a greater amount of bone in the outer portion of the defect than the interior. Additionally, the percent fill of the implants, calculated by dividing the fill fraction of the implant by the fill fraction of the surrounding bone, revealed high variability between groups, with both achieving maximum fills above 100% and minimum fills as low as 1.8% (**Table 2**). Overall, the average intensity of bone within the implants was near one, representing the bone created was similar in mineral density to the surrounding bone. There were large differences in healing, with some implants demonstrating little bone infill and others with exceptional bone infill and mineral formation. Representative micro-CT images of all constructs demonstrates that only one pig had exceptional healing in both types of implants (6365), with new bone visible between the open pores of the PCL, however, much of the PCL is still present indicated by empty spaces in the micro-CT image (**Supp. Fig. 2**). Other implants have new bone formation occurring mainly at the periphery of the implants.

**Fig. 2.**
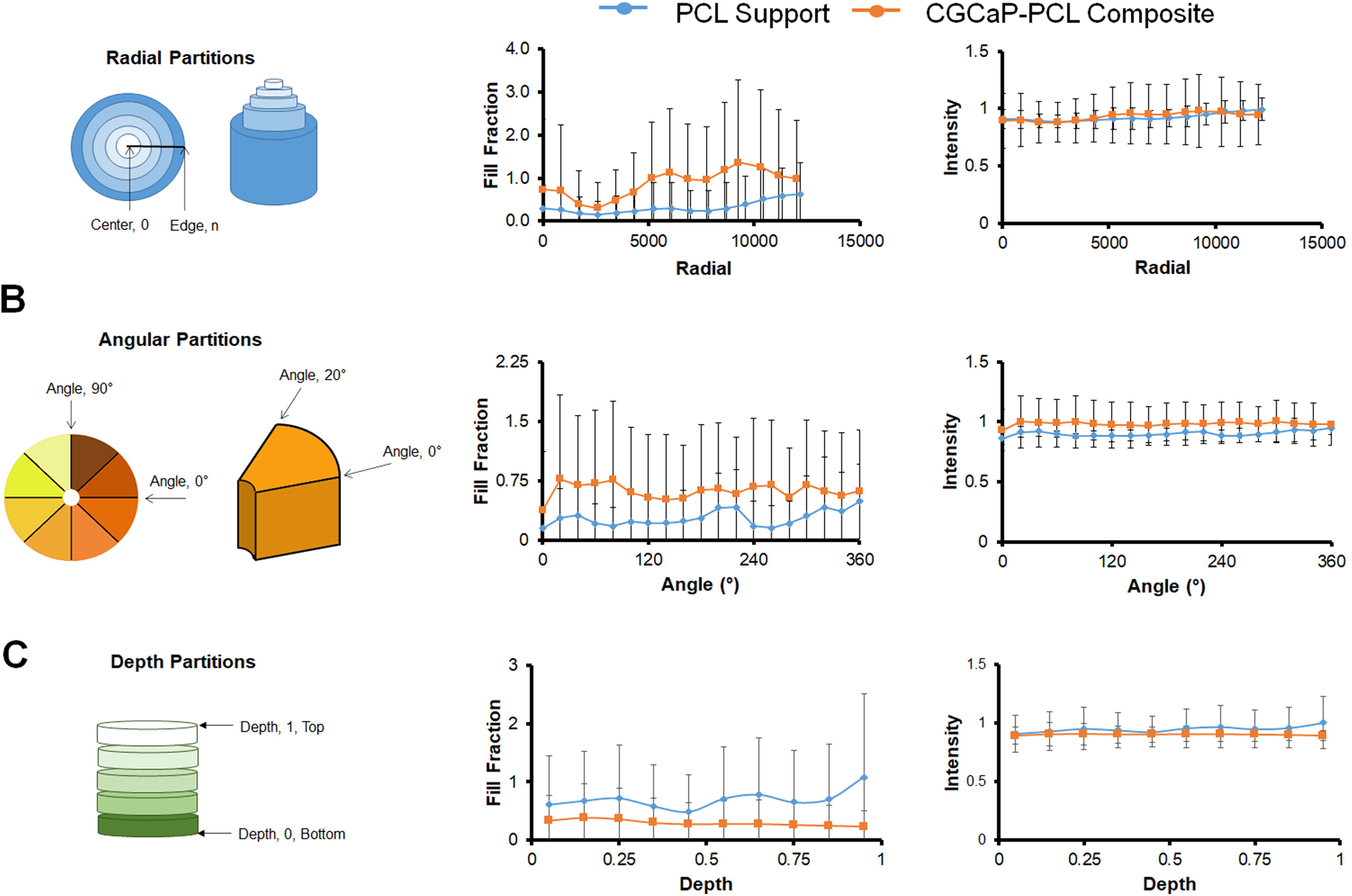
Bone infill into PCL supports and CGCaP-PCL composites as a function of the radius, angle, and depth of samples. Fill fraction values of 1 represent the same amount of bone present within the defect space as the surrounding normal bone. Intensity values of 1 represent the same mineral density present within the defect space as the surrounding normal bone. (A) Analysis of sample fill fraction and intensity by radial partitions. Center of sample is located at radial partition 0, outer edge of sample is located at approximately 12000. No significant (p < 0.05) difference between sample groups. (B) Analysis of sample fill fraction and intensity by angular partitions in 20° increments. No significant (p < 0.05) difference between sample groups. (C) Analysis of sample fill fraction and intensity by depth partitions. Bottom of sample is located at depth partition 0, top of sample is located at 1. No significant (p < 0.05) difference between sample groups. Samples were normalized to surrounding bone.

**Table 2.** Percent fill of CGCaP-PCL (Composite) compared to PCL support (PCL). The percent fill of implants was calculated by dividing the fill fraction of the implant by the fill fraction of the surrounding normal bone to get a percent fill. The implant for each pig investigated is listed.

| Fill Fraction |  |  |
| --- | --- | --- |
| Pig ID | Composite | PCL |
| 6365 | 153.6% | 100.6% |
| 6114 | 8.3% | 4.6% |
| 6363 |  | 1.8% |
| 6112 | 180.8% | 9.2% |
| 6164 |  | 30.3% |
| 6133 | 3.7% |  |
| 6375 | 4.1% |  |

### 3.3 Histology confirms limited bone formation and demonstrates fibrous tissue present

Histological examination of the implant samples confirmed results seen by micro-CT. Areas of bone tissue formation could be demonstrated in regions implicated by micro-CT (**Fig. 3**). In areas of the construct where bone tissue was not present, the PCL support were surrounded by either fibrous tissue, or fibrous tissue with significant immune infiltrate, as diagnosed by nuclear morphology (**Supp. Fig 3**). There was no discernable difference in the morphological appearance of bone tissue or of the fibrous tissue surrounding the elements of the PCL support in samples from either the CGCaP-PCL composite or the PCL supports alone. Bone tissue formed around and within the supports from both types of insert displays morphology indistinguishable from the native cancellous bone found within the jaw. Higher magnification images of the PCL support components indicated the presence of presumptive blood vessel invasion into the PCL matrix (**Supp. Fig 4**). At the periphery of the support segments, layers of fibroblast-like cells could be seen surrounding variable-sized chunks of PCL material, separating them from the body of the PCL segments (**Supp. Fig 4**). Observation of vascular infiltration and encapsulation of small portions of the PCL support material at the interface with the tissue could be demonstrated in both samples implanted with CGCaP-PCL composites and PCL supports alone, and there was no discernable difference in these observations between the two samples types.

**Fig. 3.**
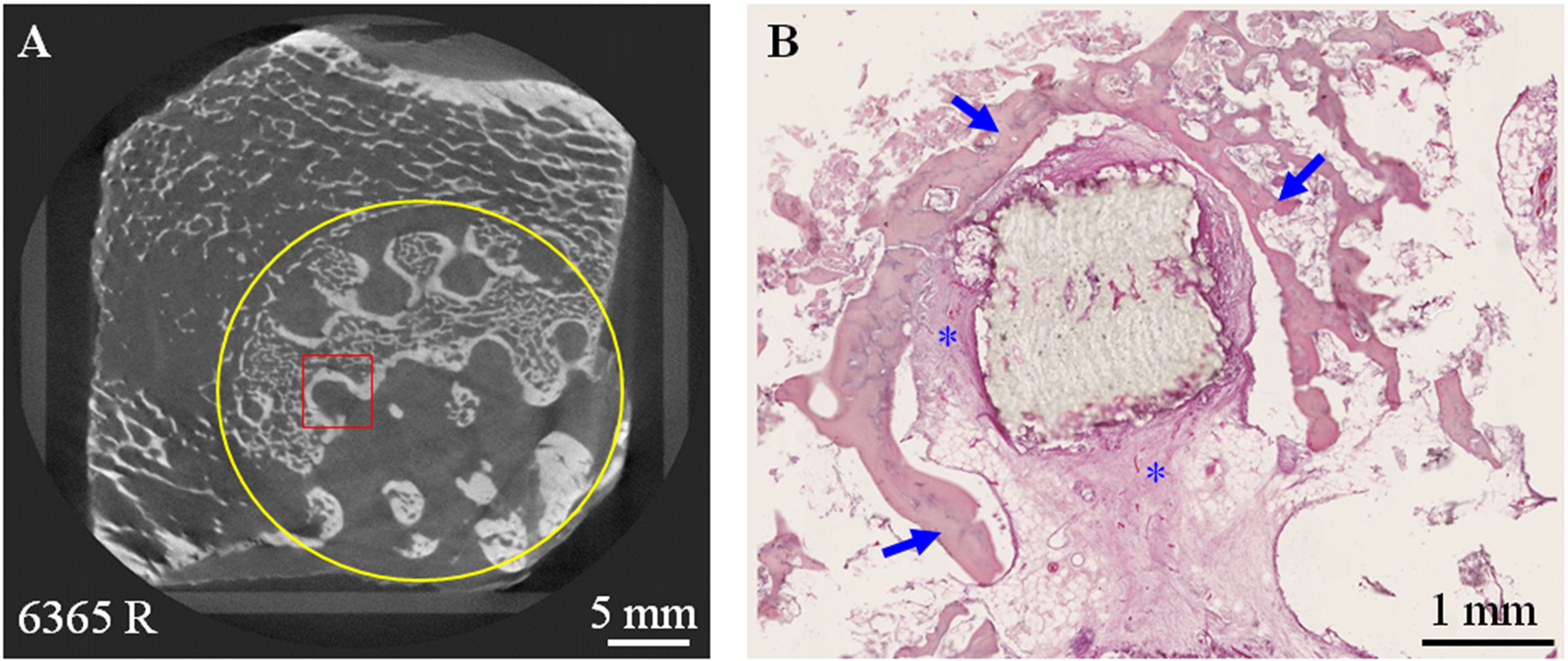
Histological confirmation of bone infill into PCL supports and CGCaP-PCL composites. (A) Micro-CT imaging of a composite insert (approximated by the yellow circle) showing mineralized tissue infiltration into the body of the composite. (B) Histological section from a region similar to the area demarcated by the red box in (A) demonstrating bone forming around a portion of PCL strut (arrows), with soft tissue forming an interface between the bone and PCL (asterisks)

## 4. Discussion

In this study, we describe the fabrication and bone regenerative capability of a mineralized collagen-PCL composite in an *in vivo* critical-sized porcine ramus defect model for CMF defect applications. CMF defect applications require a biomaterial which promotes osteogenesis and new bone formation, and in light of this study, ones which additionally promote osseointegration, fit well to the defect site, and resists chronic inflammation and bacterial infection. We have developed mineralized collagen scaffolds which promote osteogenesis and osteogenic differentiation in the absence of osteogenic supplements and promote mineral formation both *in vitro* and *in vivo* [36, 43–46]. To bolster the mechanical properties of these porous mineralized collagen scaffolds, PCL 3D-printed polymer was added to create a composite that could be made to any size or shape defect [32, 33]. Previously, the same CGCaP-PCL composites had been explored in sub-critical porcine ramus defects and had greater bone infill than mineralized collagen scaffolds or PCL supports alone, with no adverse healing [36]. However, to more accurately determine the bone regenerative effects of these CGCaP-PCL composites in CMF defect repair, the use of a critical-sized defect model was used.

To accurately evaluate healing in craniomaxillofacial defects, larger animal species are the preferred models due to their longer lifespan, immune system more closely resembling humans’, and the ability to use techniques and equipment designed for human clinical use [47]. Large animals such as cats, dogs, sheep, horses, non-human primates, and pigs are typically used to model defects and repair that could correlate to humans. Pigs in particular have similar lamellar bone structure and mineral density to humans, and results from pig studies are more likely to match studies with humans than mouse, rat, or rabbit models [47, 48]. Additionally, pigs have been shown to repair bone at approximately the same rate as humans [47]. Overall, due to the anatomy, morphology, and bone regeneration and immune response of pigs, this species is preferred to model human disease and defects [48]. The defect type must also be carefully considered for accurate translation of expected healing in humans. Sub-critical sized defects have been investigated in CMF repair, however, these defects are able to regenerate bone in the defect site over the course of the study without biomaterial intervention [49]. Critical-sized defects are large enough in size that bone will not be able to regenerate and surgical intervention is necessary to complete healing. Thus, to validate whether CGCaP-PCL composites could be a promising biomaterial in CMF defect repair, we used these in a porcine critical-sized ramus defect model.

It was not surprising to discover different results in the repair of a critical-sized defect model compared to a sub-critical sized defect model using the same biomaterial implants. Being two very different models of bone regeneration, one unable to regenerate without biomaterial assistance and the other able to fully regenerate without any implant, we can expect different healing outcomes. Here, we examined the differences in bone regeneration of a PCL support compared to a CGCaP-PCL composite. It is assumed due to the size of the defect that without an implantable material there will be no bone regeneration, thus any bone formation within the defect site is an improvement in healing. We had extremely variable healing, with one pig demonstrating bone formation throughout the defect site including highly mineralized bone, some demonstrating bone regeneration on the edges but not the center of defect space, and others only fibrous tissue formation. We had hypothesized that the CGCaP-PCL composite would form a greater amount of new bone than the PCL support alone, however, the CGCaP-PCL composites did not display significantly increased bone formation compared to the PCL supports alone, a result that did not coincide with previous sub-critical defect studies [33]. As a function of depth and angle, both CGCaP-PCL composites and PCL supports had an average bone infill less than the amount of bone surrounding the defect. However, as a function of the radius, the average bone infill increased to above 1 in the composite at the edges closest to the host bone. The overall intensity of both implants was near 1, demonstrating similar mineral intensity in new bone formed compared to surrounding bone. Additionally, percent fill of implants demonstrated a high variability in fill between groups, suggesting that while the composite approach can support significant bone regeneration, additional innovations are required to promote consistent responses. It is notable that while the 25 mm diameter critical sized defect used here does not show any innate healing capacity [50], both PCL and CGCaP-PCL implants were able to induce quantifiable, and some cases signficnt new bone regeneration with the quantity and quality of regenerated bone showing the potential to match the native surrounding bone. One pig in particular demonstrated new bone formation throughout the pores of the PCL support and CGCaP-PCL composite, extending new bone formation to the center of the defect space. Where new bone was present, it surrounded the PCL support structures, with an interface of soft/fibrous tissue between the surface of the bone and the surface of the PCL (**Fig. 3**). The soft/fibrous tissue always made contact with the PCL, not the bone itself directly. This soft tissue appeared to penetrate the surface of the PCL, carving channels in it; allowing penetration of the PCL with presumptive vasculature, and separating small pieces of the PCL off from the main body of the PCL structure at the periphery (**Supp. Fig. 4**). This histology was similar in both PCL implants and CGCaP-PCL composite implants.

Bone infill into the implant was limited by the PCL polymer, which can be visualized in **Supp. Fig. 2**, with gaps in the bone formation from PCL still present. Due to the long degradation time of PCL (2+ years), there was still a significant amount of PCL left in the wound site. Fibrous tissue formation could also be attributed to the degradation byproducts of PCL and the size and shape of the print. The size and shape of polymer implants have been related to fibrous tissue formation and more PCL present results in more acidic byproduct release during degradation [51, 52]. Issues in healing could also be attributed to the fit of the polymer and the composite to the defect site. Five implants were visibly ejected from the jaws of the pig or were unable to be found during micro-CT, which could be due to the method of press-fitting and gluing scaffolds and composites into the defect site. Unstable implant fixation and micromotion of implants has been shown to produce a fibrous tissue response. Micromotion resulting from unstable fixation of the implants could be a potential explanation for this tissue formation within both PCL supports and CGCaP-PCL composites [10, 11].

Multiple implants, both PCL supports and CGCaP-PCL composites, had post-surgical complications. Two pigs suffered from abscess formation, possibly due to bacterial infection during open wound fractures [16]. Although antibiotics were used during surgery, bacteria associated with infection in bone have demonstrated resistance and have been linked to disrupting osteoblast activity and bone formation [17]. In most of these implants, little bone was formed, and histology of samples with little to no bone fill frequently showed significant leukocyte infiltration in the fibrous tissue surrounding the PCL supports (**Supp. Fig. 3**). This suggests chronic inflammation or presence of infection, and this could potentially explain the lack of bone tissue formation in these samples, which could potentially be avoided by more rigorous sterilization of medical tools and devices.

It should be noted that out of 22 implanted materials, 2 of these had exceptional healing outcomes. Both the CGCaP-PCL and PCL support within one pig demonstrated bone infill throughout the entirety of the implant and integration with surrounding host bone. Additionally, this implant did not have any abscess or fibrous tissue formation. Although other implants had difficulty healing throughout the entire space, successful healing and bridged bone formation from the center to the periphery of the defect in one implant demonstrates the potential for this strategy to work to regenerate bone in critical-sized defects. New bone formation would not have occurred without biomaterial intervention, and even small amounts of new bone formation in multiple implants demonstrate the potential for this strategy to succeed if the challenges in CMF defect repair are addressed. This study therefore identifies a series of modifications to address observed challenges in repair.

The limited bone formation and fibrous tissue encapsulation could be attributed to the known obstacles of CMF defect repair: implant fit to host bone, chronic inflammation and infection, and promoting osteogenesis. Due to variable healing results seen with micro-CT and histology, we recognize the need for improved design of these implants. The PCL polymer present in our implants was printed in a design that did not allow for any shape-fitting, which can be used to minimize micromotion. Micromotion ultimately can inhibit osseointegration of the implant and surrounding bone [10, 11]. In the future we plan to modify the polymer 3D-print design to allow for shape-fitting, as well as minimize the amount and type of polymer used to shorten the degradation time and allow for greater bone infill. We have already demonstrated that we can incorporate other 3D-printed structures into these collagen scaffolds to promote greater mineral formation and faster degradation times [53, 54]. There were multiple abscesses and fibrous tissue formed, which can be related to bacterial infection and the body’s natural instinct towards inflammation during healing. Due to the large size of bone missing, this makes bacterial infection more likely, as well as persistent inflammation, and a biomaterial that can prevent infection and guide the immune response towards repair should lead to more successful bone repair [14, 15, 55]. Future efforts are focusing on changes to scaffold compositional elements to modulate immune cell phenotype [56, 57]. Finally, limited bone formation in many of the implants, regardless of implant type, was different from the performance of these same groups in sub-critical sized defects. The mineralized collagen scaffold robustly promotes osteogenesis *in vitro* and in small animal studies *in vivo* [20, 21, 29–31]. However, the large scale of the defects tested likely require efforts to boost cell recruitment and activity. Key modification include: the use of intraoperative adipose-derived stem cells seeding to boost healing [58]; modifications to the glycosaminoglycan content and pore size or orientation of the mineralized collagen scaffold to promote greater cell migration into the defect [57, 59] and the potential to incorporate biomolecular signals (e.g., transiently incorporated BMP2, VEGF) or modify the mineral composition (e.g., inclusion of zinc ions) to promote proliferation, osteogenesis, or angiogenic activity [60–62].

Our investigative team has demonstrated the individual potential of each of these interventions via *in vitro* or *in vivo* studies, suggesting routes for future improvements of these collagen-PCL composites to meet the translational requirements of regenerative healing of critical-sized craniofacial bone defects.

## 5. Conclusions

The bone regenerative capacity of PCL 3D-printed supports and mineralized collagen combined with PCL supports was investigated in a critical-sized porcine ramus defect model. We found variable healing, with the average bone infill within the implants to be less than the amount of bone surrounding these, however, greater than the amount of new bone formation possible without these implants. One implant demonstrated new bone formation throughout the space of the defect, and others demonstrated new bone formation mainly at the edges of the defect space. Some implants displayed fibrous tissue formation preventing new bone formation, possibly due to the amount of polymer present, infection, and lack of implant stability within the defect space. While there were no significant differences in healing between the PCL supports and CGCaP-PCL composites, observations strongly motivate future changes in composite biomaterials to improve regenerative healing of large-scale craniofacial bone defects. The instance of bone formation throughout the defect space in both implants demonstrates the potential for this strategy to work with further modifications. This study motivates future work to improve cell invasion and resultant osteogenesis, improve conformal fitting to minimize micromotion and resultant fibrous tissue formation, and modulate the immune response to aid in repair.

## Acknowledgements

The authors would like to acknowledge the University of Illinois the Carl R. Woese Institute for Genomic Biology, the Chemical and Biomolecular Engineering Department, the Imported Swine Research Laboratory and the Beckman Institute for Advanced Science and Technology, all located at the University of Illinois at Urbana-Champaign. The authors would also like to thank Leilei Yin in the Imaging Technology Group at Beckman Institute for assistance with Micro-CT. The authors would also like to acknowledge Eileen Johnson and Simona Slater for assistance in data analysis. Research reported in this publication was supported by the AO Foundation (Switzerland) as Project S-14-54H. Research reported in this publication was also supported by the National Institute of Dental and Craniofacial Research of the National Institutes of Health under Award Number R21 DE026582. The content is solely the responsibility of the authors and does not necessarily represent the official views of the NIH. We are grateful for the funding for this study provided by the NSF Graduate Research Fellowship DGE-1144245 (MD).

## Ethical Statement

All animal experimentation protocols were approved by the University of Illinois at Urbana-Champaign Institutional Animal Care and Use Committee under protocol #15065.

## References

[1] E.C. Kruijt Spanjer, G.K.P. Bittermann, I.E.M. van Hooijdonk, A.J.W.P. Rosenberg, D. Gawlitta, Taking the endochondral route to craniomaxillofacial bone regeneration: A logical approach?, Journal of Cranio-Maxillofacial Surgery 45 (2017) 1099–1106.

[2] A. Norozy, M.H.K. Motamedi, A. Ebrahimi, H. Khoshmohabat, Maxillofacial Fracture Patterns in Military Casualties, Journal of Oral and Maxillofacial Surgery 78(4) (2020) 611.e1–611.e6.

[3] A. Depeyre, S. Touzet-Roumazeille, L. Lauwers, G. Raoul, J. Ferri, Retrospective evaluation of 211 patients with maxillofacial reconstruction using parietal bone graft for implants insertion, Journal of Cranio-Maxillofacial Surgery 44 (2016) 1162–1169.

[4] A. Abuzayed, S. Aydin, S. Aydin, B. Kucukyuruk, G. Sanus, Cranioplasty: Review of materials and techniques, Journal of Neurosciences in Rural Practice 2 (2011) 162.

[5] M. Elsalanty, D. Genecov, Bone Grafts in Craniofacial Surgery, Craniomaxillofacial Trauma and Reconstruction 2 (2009) 125–134.

[6] S. Bose, M. Roy, A. Bandyopadhyay, Recent advances in bone tissue engineering scaffolds, Trends Biotechnol. 30 (2012) 546–554.

[7] W. Wang, K.W.K. Yeung, Bone grafts and biomaterials substitutes for bone defect repair: A review, Bioactive Materials 2 (2017) 224–247.

[8] K.A. Hing, Bone repair in the twenty-first century: Biology, chemistry or engineering?, Philosophical Transactions of the Royal Society A: Mathematical, Physical and Engineering Sciences 362 (2004) 2821–2850.

[9] E. Gibon, L.Y. Lu, K. Nathan, S.B. Goodman, Inflammation, ageing, and bone regeneration, Journal of Orthopaedic Translation 10 (2017) 28–35.

[10] M.J. Cross, G.J. Roger, J. Spycher, Cementless fixation techniques and challenges in joint replacement, in: P.A. Revell (Ed.), Joint Replacement Technology, Woodhead Publishing Limited, Cambridge, UK, 2014, pp. 186–211.

[11] R. Dimitriou, G.C. Babis, Biomaterial osseointegration enhancement with biophysical stimulation, Journal of Musculoskeletal Neuronal Interactions 7 (2007) 253–265.

[12] R.A. Hortensius, B.A.C. Harley, Naturally derived biomaterials for addressing inflammation in tissue regeneration, Experimental Biology and Medicine 241 (2016) 1015–1024.

[13] T. Wynn, T. Ramalingam, Mechanisms of fibrosis: therapeutic translation for fibrotic disease, Nature Medicine 18 (2013) 1028–1040.

[14] J.M. Anderson, Inflammation, Wound Healing, and the Foreign-Body Response, in: B. Ratner, A. Hoffman, F.J. Schoen, J. Lemons (Eds.), Biomaterials Science, Academic Press, United States, 1996, pp. 503–512.

[15] M. Ribeiro, F.J. Monteiro, M.P. Ferraz, Infection of orthopedic implants with emphasis on bacterial adhesion process and techniques used in studying bacterial-material interactions., Biomatter 2 (2012) 176–194.

[16] A. Trampuz, W. Zimmerli, Diagnosis and treatment of implant-associated septic arthritis and osteomyelitis, Current Infectious Disease Reports 10 (2008) 394–403.

[17] J. Josse, F. Velard, S. Gangloff, Staphylococcus aureus vs. Osteoblast: Relationship and Consequences in Osteomyelitis, Front Cell Infect Microbiol 5(85) (2015) 1–17.

[18] A. Weisgerber, S. Caliari, B. Harley, Mineralized collagen scaffolds induce hMSC osteogenesis and matrix remodeling, Biomaterials Science 3 (2015) 533–42.

[19] A. Farrell, I. Yannas, B. O’Connell, P.J. Prendergast, V.A. Campbell, P. Doyle, B.A. Harley, F.J. O’Brien, J. Fischer, A Collagen-glycosaminoglycan Scaffold Supports Adult Rat Mesenchymal Stem Cell Differentiation Along Osteogenic and Chondrogenic Routes, Tissue Engineering 12 (2006) 459–468.

[20] X. Ren, D.W. Weisgerber, D. Bischoff, M.S. Lewis, R.R. Reid, T.C. He, D.T. Yamaguchi, T.A. Miller, B.A.C. Harley, J.C. Lee, Nanoparticulate Mineralized Collagen Scaffolds and BMP-9 Induce a Long-Term Bone Cartilage Construct in Human Mesenchymal Stem Cells, Advanced Healthcare Materials 5 (2016) 1821–1830.

[21] X. Ren, D. Bischoff, D.W. Weisgerber, M.S. Lewis, V. Tu, D.T. Yamaguchi, T.A. Miller, B.A.C. Harley, J.C. Lee, Osteogenesis on nanoparticulate mineralized collagen scaffolds via autogenous activation of the canonical BMP receptor signaling pathway, Biomaterials 50 (2015) 107–114.

[22] Y.-j. Seong, I.-g. Kang, E.-h. Song, H.-e. Kim, S.-h. Jeong, Calcium Phosphate – Collagen Scaffold with Aligned Pore Channels for Enhanced Osteochondral Regeneration, Advanced Healthcare Materials 6 (2017) 1–11.

[23] J.P. Gleeson, T. Weber, T. Levingstone, A.A. Al-Munajjed, F.J. O’Brien, J. Hammer, N.A. Plunkett, C. Jungreuthmayer, Development of a biomimetic collagen-hydroxyapatite scaffold for bone tissue engineering using a SBF immersion technique, Journal of Biomedical Materials Research Part B: Applied Biomaterials 90B (2009) 584–591.

[24] A. Al-Munajjed, J. Gleeson, F. O’Brien, Development of a collagen calcium-phosphate scaffold as a novel bone graft substitute., Stud Health Technol Inform 133 (2008) 11–20.

[25] M. Tang, W. Chen, J. Liu, M.D. Weir, L. Cheng, H.H. Xu, Human induced pluripotent stem cell-derived mesenchymal stem cell seeding on calcium phosphate scaffold for bone regeneration, Tissue Eng Part A 20(7–8) (2014) 1295–305.

[26] A. Zaopo, R. Papagna, P. Mimo, F. Carini, E. Donzelli, A. Salvadè, M. Miloso, M. Baldoni, M. Morrone, G. Tredici, M. Viganò, Mesenchymal stem cells cultured on a collagen scaffold: In vitro osteogenic differentiation, Archives of Oral Biology 52 (2006) 64–73.

[27] G.S. Offeddu, J.C. Ashworth, R.E. Cameron, M.L. Oyen, Multi-scale mechanical response of freeze-dried collagen scaffolds for tissue engineering applications, Journal of the Mechanical Behavior of Biomedical Materials 42 (2015) 19–25.

[28] F.G. Lyons, J.P. Gleeson, S. Partap, K. Coghlan, F.J. O’Brien, Novel microhydroxyapatite particles in a collagen scaffold: a bioactive bone void filler?, Clinical Orthopaedics and Related Research 472 (2014) 1318–28.

[29] X. Ren, Q. Zhou, D. Foulad, M.J. Dewey, D. Bischoff, T.A. Miller, D.T. Yamaguchi, B.A.C. Harley, J.C. Lee, Nanoparticulate mineralized collagen glycosaminoglycan materials directly and indirectly inhibit osteoclastogenesis and osteoclast activation, Journal of tissue engineering and regenerative medicine 13(5) (2019) 823–834.

[30] X. Ren, Q. Zhou, D. Foulad, A.S. Tiffany, M.J. Dewey, D. Bischoff, T.A. Miller, R.R. Reid, T.-c. He, D.T. Yamaguchi, B.A.C. Harley, J.C. Lee, Osteoprotegerin reduces osteoclast resorption activity without affecting osteogenesis on nanoparticulate mineralized collagen scaffolds, Science Advances 5 (2019) 1–12.

[31] X. Ren, V. Tu, D. Bischoff, D.W. Weisgerber, M.S. Lewis, D.T. Yamaguchi, T.A. Miller, B.A. Harley, J.C. Lee, Nanoparticulate mineralized collagen scaffolds induce in vivo bone regeneration independent of progenitor cell loading or exogenous growth factor stimulation, Biomaterials 89 (2016) 67–78.

[32] D.W. Weisgerber, K. Erning, C.L. Flanagan, S.J. Hollister, B.A.C. Harley, Evaluation of multi-scale mineralized collagen-polycaprolactone composites for bone tissue engineering, Journal of the Mechanical Behavior of Biomedical Materials 61 (2016) 318–327.

[33] A. Weisgerber, D. Milner, H. Lopez-Lake, M. Rubessa, S. Lotti, K. Polkoff, R. Hortensius, C. Flanagan, S. Hollister, M. Wheeler, B. Harley, A mineralized collagen-polycaprolactone composite promotes healing of a porcine mandibular ramus defect, Tissue Eng Part A 0 (2017) 1–12.

[34] S. Hollister, C. Lin, E. Saito, C. Lin, R. Schek, J. Taboas, J. Williams, B. Partee, C. Flanagan, A. Diggs, E. Wilke, G.V. Lenthe, R. Muller, T. Wirtz, S. Das, S. Feinberg, P. Krebsbach, Engineering craniofacial scaffolds, Orthod Craniofacial Res 8 (2005) 162–173.

[35] A.G. Mitsak, J.M. Kemppainen, M.T. Harris, S.J. Hollister, Effect of polycaprolactone scaffold permeability on bone regeneration in vivo, Tissue Eng Part A 17(13–14) (2011) 1831–9.

[36] D.W. Weisgerber, D.J. Milner, H. Lopez-Lake, M. Rubessa, S. Lotti, K. Polkoff, R.A. Hortensius, C.L. Flanagan, S.J. Hollister, M.B. Wheeler, B.A.C. Harley, A Mineralized Collagen-Polycaprolactone Composite Promotes Healing of a Porcine Mandibular Defect, Tissue Eng Part A 24(11–12) (2018) 943–954.

[37] D.W. Weisgerber, S.R. Caliari, B.A.C. Harley, Mineralized collagen scaffolds induce hMSC osteogenesis and matrix remodeling, Biomater Sci 3(3) (2015) 533–42.

[38] S.R. Caliari, B.A.C. Harley, Structural and biochemical modification of a collagen scaffold to selectively enhance MSC tenogenic, chondrogenic, and osteogenic differentiation, Advanced healthcare materials 3(7) (2014) 1086–96.

[39] S.R. Caliari, B.A.C. Harley, The effect of anisotropic collagen-GAG scaffolds and growth factor supplementation on tendon cell recruitment, alignment, and metabolic activity, Biomaterials 32(23) (2011) 5330–40.

[40] L.H.H. Olde Damink, P.J. Dijkstra, M.J.A. van Luyn, P.B. Van Wachem, P. Nieuwenhuis, J. Feijen, Cross-linking of dermal sheep collagen using a water soluble carbodiimide, Biomaterials 17 (1996) 765–773.

[41] N.H. Veilleux, I.V. Yannas, M. Spector, Effect of passage number and collagen type on the proliferative, biosynthetic, and contractile activity of adult canine articular chondrocytes in type I and II collagen-glycosaminoglycan matrices in vitro, Tissue Eng 10(1–2) (2004) 119–27.

[42] R.L.O.a.L. Necker, An Introduction to Statistical Methods and Data Analysis, 7 ed., Cengage Learning2010.

[43] D.W. Weisgerber, S.R. Caliari, B.A. Harley, Mineralized collagen scaffolds induce hMSC osteogenesis and matrix remodeling, Biomater Sci 3(3) (2015) 533–42.

[44] S.R. Caliari, B.A. Harley, Structural and biochemical modification of a collagen scaffold to selectively enhance MSC tenogenic, chondrogenic, and osteogenic differentiation, Adv Healthc Mater 3(7) (2014) 1086–96.

[45] X. Ren, D. Bischoff, D.W. Weisgerber, M.S. Lewis, V. Tu, D.T. Yamaguchi, T.A. Miller, B.A. Harley, J.C. Lee, Osteogenesis on nanoparticulate mineralized collagen scaffolds via autogenous activation of the canonical BMP receptor signaling pathway, Biomaterials 50 (2015) 107–14.

[46] X. Ren, D.W. Weisgerber, D. Bischoff, M.S. Lewis, R.R. Reid, T.C. He, D.T. Yamaguchi, T.A. Miller, B.A. Harley, J.C. Lee, Nanoparticulate Mineralized Collagen Scaffolds and BMP-9 Induce a Long-Term Bone Cartilage Construct in Human Mesenchymal Stem Cells, Adv Healthc Mater 5(14) (2016) 1821–30.

[47] M. Rubessa, K. Polkoff, M. Bionaz, E. Monaco, D.J. Milner, S.J. Holllister, M.S. Goldwasser, M.B. Wheeler, Use of Pig as a Model for Mesenchymal Stem Cell Therapies for Bone Regeneration, Anim Biotechnol 28(4) (2017) 275–287.

[48] J. Stembirek, M. Kyllar, I. Putnova, L. Stehlik, M. Buchtova, The pig as an experimental model for clinical craniofacial research, Lab Anim 46(4) (2012) 269–79.

[49] A. Gosain, L. Song, P. Yu, B. Mehrara, C. Maeda, L. Gold, M. Longaker, Osteogenesis in Cranial Defects: Reassessment of the Concept of Critical Size and the Expression of TGF-b Isoforms, Plastic and Reconstructive Surgery 106(2) (2000) 360–371.

[50] S.M. Wilson, M.S. Goldwasser, S.G. Clark, E. Monaco, M. Bionaz, W.L. Hurley, S. Rodriguez-Zas, L. Feng, Z. Dymon, M.B. Wheeler, Adipose-derived mesenchymal stem cells enhance healing of mandibular defects in the ramus of swine, Journal of Oral and Maxillofacial Surgery 70 (2012) e193–e203.

[51] N.L. Davison, F. Barrère-de Groot, D.W. Grijpma, Degradation of Biomaterials, in: C.A.V. Blitterswijk, J.D. Boer (Eds.), Tissue Engineering, Academic Press, United States, 2015, pp. 177–215.

[52] M.A. Woodruff, D.W. Hutmacher, The return of a forgotten polymer – Polycaprolactone in the 21st century, Progress in Polymer Science (Oxford) 35 (2010) 1217–1256.

[53] M.J. Dewey, E.M. Johnson, D.W. Weisgerber, M.B. Wheeler, B.A.C. Harley, Shape-fitting collagen-PLA composite promotes osteogenic differentiation of porcine adipose stem cells, Journal of the Mechanical Behavior of Biomedical Materials 95 (2019) 21–33.

[54] M.J. Dewey, A.V. Nosatov, K. Subedi, R. Shah, A. Jakus, B.A.C. Harley, Inclusion of a 3D-printed Hyperelastic Bone mesh improves mechanical and osteogenic performance of a mineralized collagen scaffold, Acta Biomaterialia 121 (2021) 224–236.

[55] J.M. Anderson, Foreign body reaction to biometarials, Semin Immunol. 20 (2008) 86–100.

[56] H. Niknejad, H. Peirovi, A. Seifalian, A. Ahmadiani, J. Ghanavi, M. Jorjani, Properties of the amniotic membrane for potential use in tissue engineering, European Cells and Materials 15 (2008) 88–99.

[57] M.J. Dewey, E.M. Johnson, S.T. Slater, D.J. Milner, M.B. Wheeler, B.A.C. Harley, Mineralized collagen scaffolds fabricated with amniotic membrane matrix increase osteogenesis under inflammatory conditions, Regenerative Biomaterials rbaa005 (2020) 1–12.

[58] S.M. Wilson, M.S. Goldwasser, S.G. Clark, E. Monaco, M. Bionaz, W.L. Hurley, S. Rodriguez-Zas, L. Feng, Z. Dymon, M.B. Wheeler, Adipose-derived mesenchymal stem cells enhance healing of mandibular defects in the ramus of swine, J Oral Maxillofac Surg 70(3) (2012) e193–203.

[59] M.J. Dewey, A.V. Nosatov, K. Subedi, B. Harley, Anisotropic mineralized collagen scaffolds accelerate osteogenic response in a glycosaminoglycan-dependent fashion, RSC Advances 10(26) (2020) 15629–15641.

[60] A.S. Tiffany, M.J. Dewey, B.A.C. Harley, Sequential sequestrations increase the incorporation and retention of multiple growth factors in mineralized collagen scaffolds, RSC Adv 10 (2020) 26982–96.

[61] A.S. Tiffany, D.L. Gray, T.J. Woods, K. Subedi, B.A.C. Harley, The inclusion of zinc into mineralized collagen scaffolds for craniofacial bone repair applications, Acta Biomaterialia 93 (2019) 86–96.

[62] W.K. Grier, A.S. Tiffany, M.D. Ramsey, B.A.C. Harley, Incorporating β-cyclodextrin into collagen scaffolds to sequester growth factors and modulate mesenchymal stem cell activity, Acta Biomaterialia 76 (2018) 116–125.

