## Supplementary Files for "Repair of critical-size porcine craniofacial bone defects using a collagen-polycaprolactone composite biomaterial"

### **Supplementary Figures**

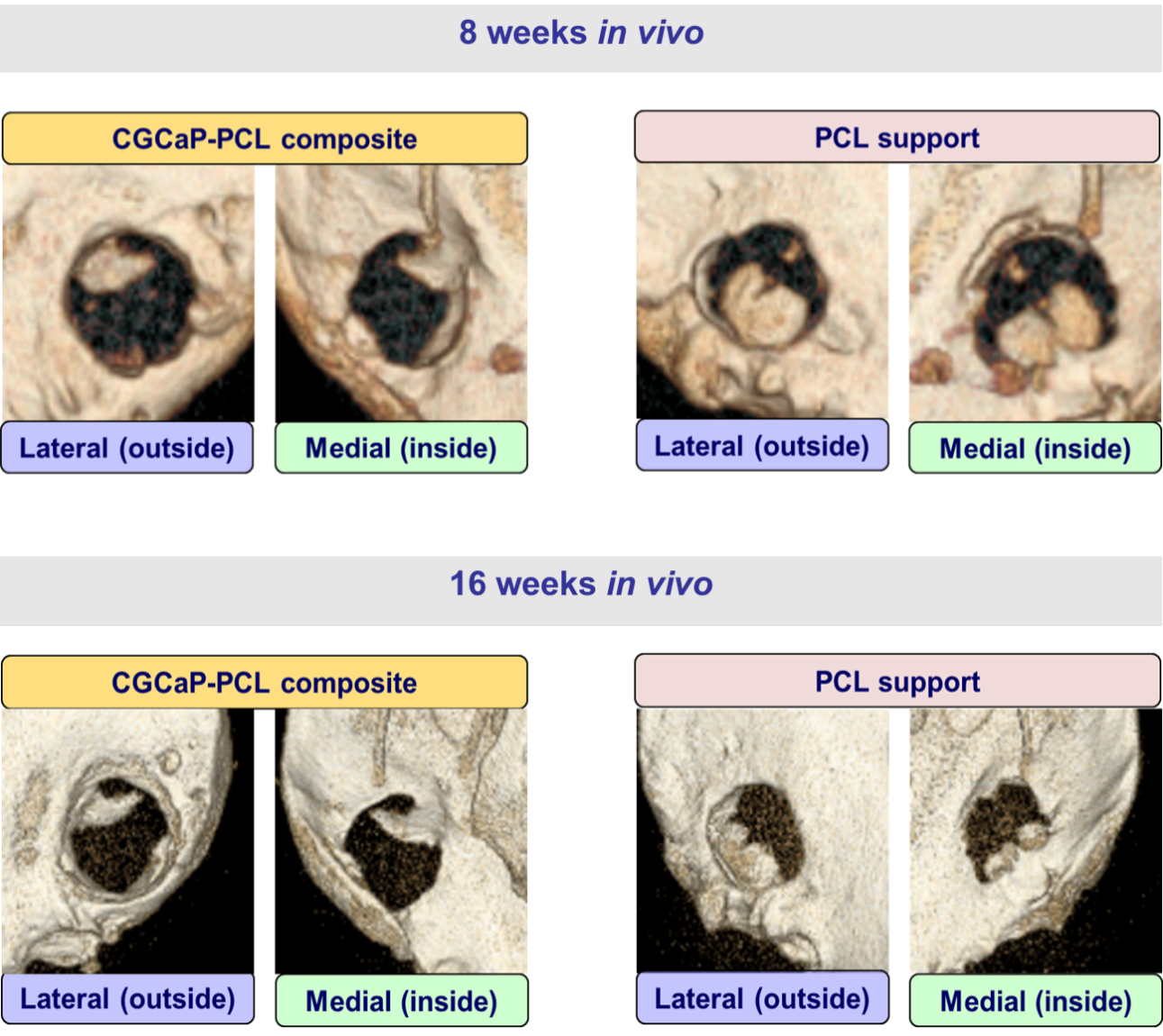

**Supplementary Figure 1.** **Micro-CT images of live pigs at 8 weeks and 16 weeks with PCL supports and CGCaP-PCL composites.**

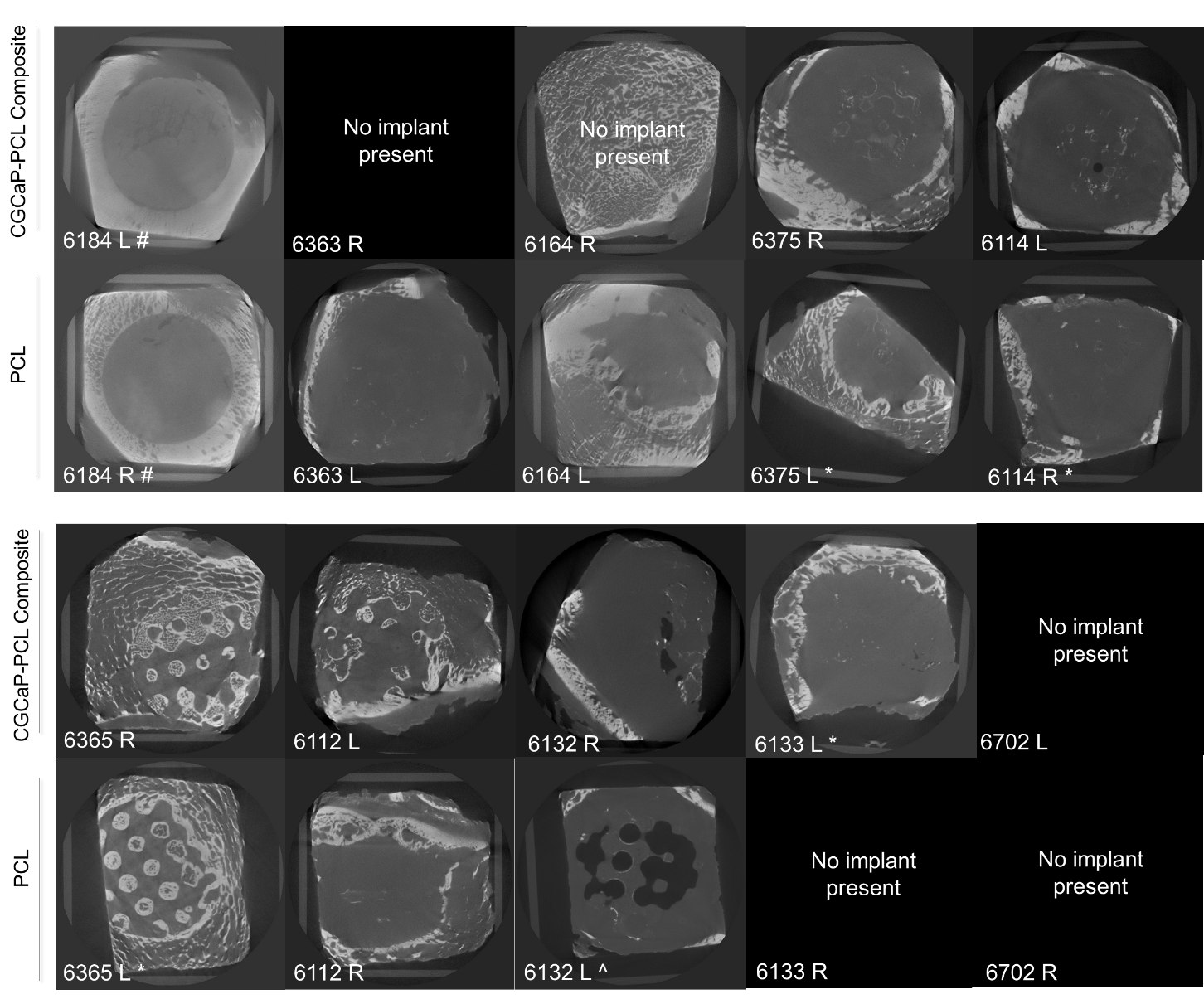

**Supplementary Figure 2. Representative Micro-CT images of CGCaP-PCL Composite and PCL support in the left and right mandibles of pigs after 9-10 months of implantation.** # represents pig was sacrificed early due to health issues (6 days post implantation). ‘No implant present’ indicates that the CGCaP-PCL composite or PCL support was not found in the pig mandible or migrated out of the mandible during healing. * represents implant had an edge partially cut off during trimming procedure. ^ represents abscess was present over the implant.

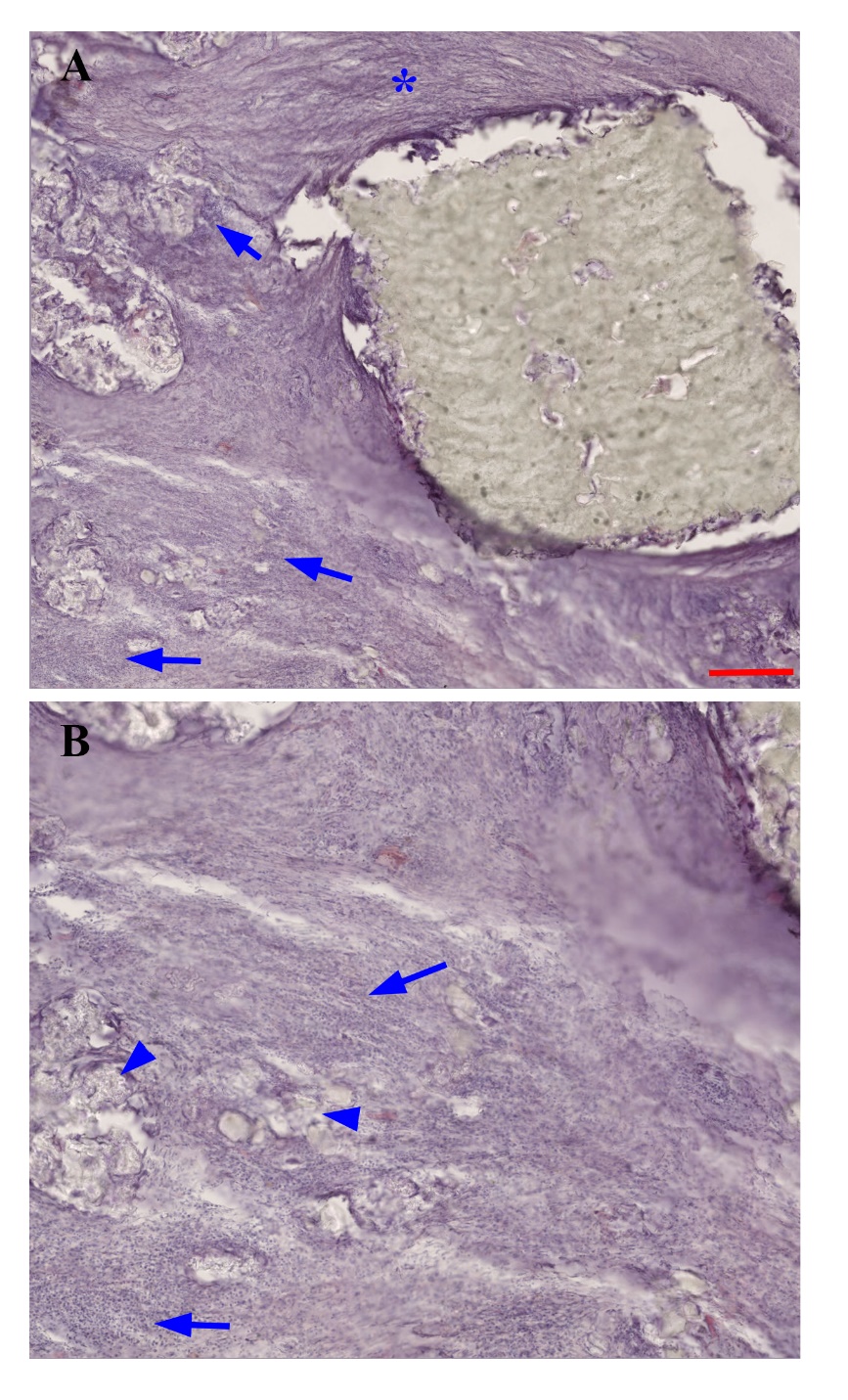

**Supplementary Figure 3. Encapsulation of PCL Strut Forms in Fibrous Tissue and Infiltrating Immune cells.** (A) Hematoxylin and eosin stained section of a PCL strut surrounded by fibrous connective tissue (asterisk) and significant infiltration of immune cells (arrows). Scale bar: 500 μm. (B) Enlarged region from (A) showing clusters of infiltrating lymphocytes (arrows). Note the presence of small shards of polycaprolactone embedded in these cell clusters.

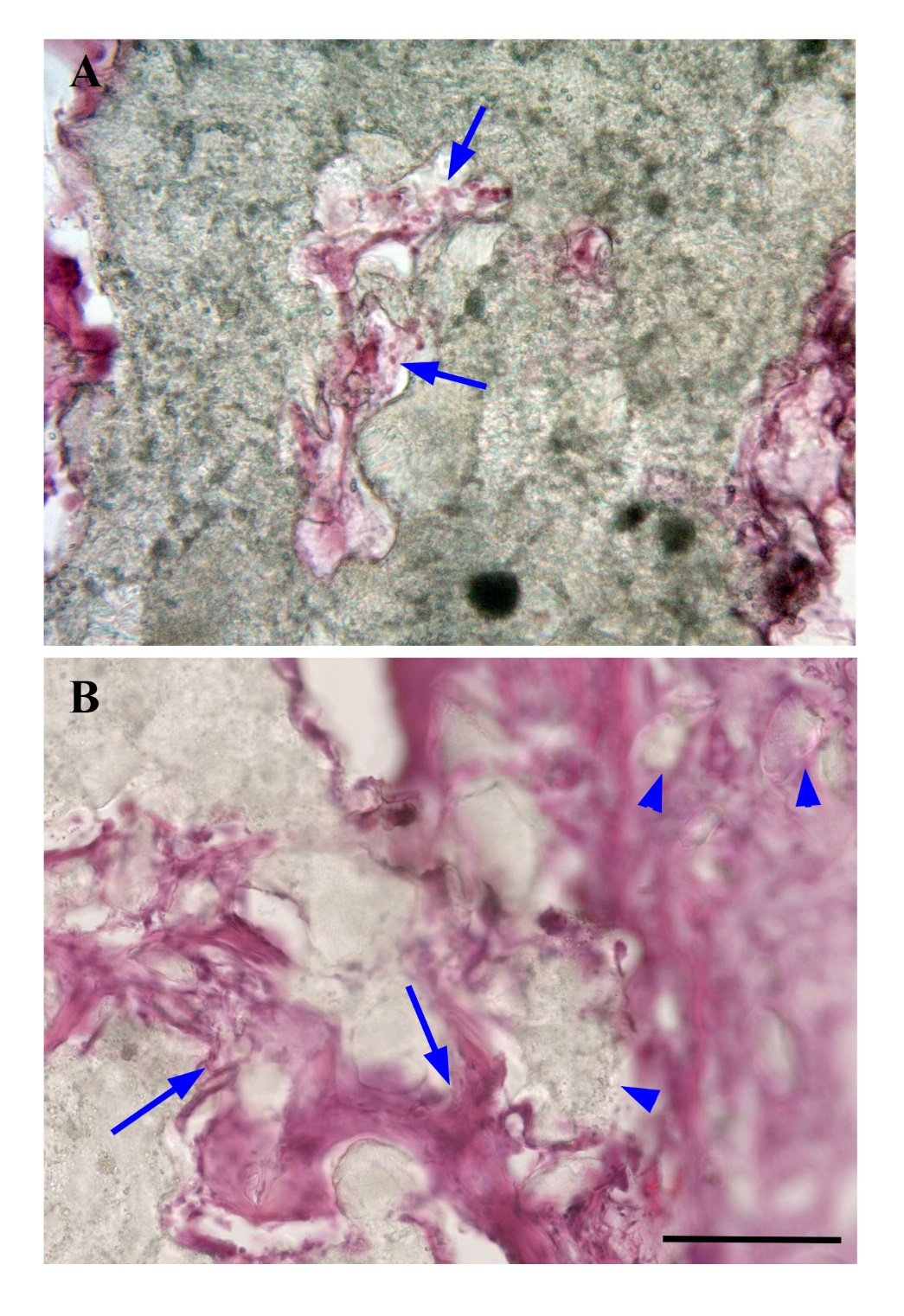

**Supplementary Figure 4. Invasion of Polycaprolactone Support Structures by Vasculature and Connective Tissue.** (A) Penetration of PCL matrix by vascular tissue (presumptive capillaries) as diagnosed by the presence of red blood cells (arrows) in these vessel-like structures. (B) Penetration of the periphery of a PCL support strut with vasculature and connective tissue (arrows). As the tissue proceeds inward, small pieces of polycaprolactone are surrounded by connective tissue and pinched off from the main body of the polycaprolactone strut (arrowheads). Penetration of PCL struts is observed both in samples surrounded by bone ingrowth, and samples surrounded by fibrous tissue and immune infiltrates. Scale bar: 100 μm.
